# Reconstructing N-Glycan Profiles of Individual Glycoproteins from the Total Blood Plasma N-Glycome: Implications for Immunoglobulin G N-Glycosylation GWAS

**DOI:** 10.64898/2026.01.30.702519

**Authors:** Anna Soplenkova, Denis Maslov, Anna Timoshchuk, Domagoj Kifer, Ana Cvetko, Michel Georges, Claire J. Steves, Cristina Menni, Sodbo Sharapov, Gordan Lauc, Yurii S. Aulchenko

## Abstract

The genetic regulation of the plasma N-glycome variation in human populations is not fully characterized, partly due to the limited sample size in glyco-genetics studies. Here, we aimed to demonstrate that protein-specific N-glycan profiles, like those of immunoglobulin G (IgG), can be accurately reconstructed from the total plasma N-glycome (TPNG), enabling us to find new regulators of this complex process re-analysing existing datasets. By testing multiple linear and non-linear machine learning approaches we built a model to reconstruct IgG N-glycans from TPNG data, training on the TwinsUK cohort and validating on CEDAR. We reconstruct GWAS summary statistics for IgG N-glycans by applying the trained linear model to plasma glycan GWAS summary statistics, i.e., as GWAS of linear combinations of plasma glycan traits. The majority of the identified loci had been implicated in IgG N-glycosylation GWAS. Additionally, we found four new loci and suggested the role of FCRLA, KDELR2, HHEX, and TCF3 in the regulation of IgG N-glycosylation. In conclusion, we showed that our method enables the creation of protein-specific N-glycome datasets, allowing for powerful meta-analyses without the need to profile new samples.

## Introduction

Glycosylation is a common post- and co-translational modification of proteins involving the enzymatic addition of saccharide chains, glycans, to proteins^1^. This process significantly influences both the structure and the function of proteins^2–5^. As a highly regulated process, glycosylation produces various glycoforms that contribute to different biological properties^6^. The synthesis of glycans primarily occurs in the endoplasmic reticulum and Golgi apparatus, facilitated by the coordinated actions of glycosyltransferases and other enzymes that manage the availability of substrates essential for glycan assembly^7^ and their regulators.

While the biochemical network of human N-glycan biosynthesis is well understood^8^, little is known about *in vivo* regulation of this process^9^, including tissue- and protein-specific regulation. Immunoglobulins, which are synthesized by antibody-producing B-cells, and glycoproteins produced by hepatocytes in the liver form the majority of plasma glycoproteins^10^ (Figure 1a). As immunoglobulin G (IgG) is the most abundant glycoprotein in total blood plasma, IgG N-glycome and total plasma N-glycome (TPNG) are quite strongly correlated with each other (Supplementary Figure 1). This suggests that IgG N-glycome can be reconstructed from plasma N-glycome. However, an accuracy of reconstruction is obscured by the complex mixture of other N-glycosylated proteins present in human blood plasma.

**Figure 1:**
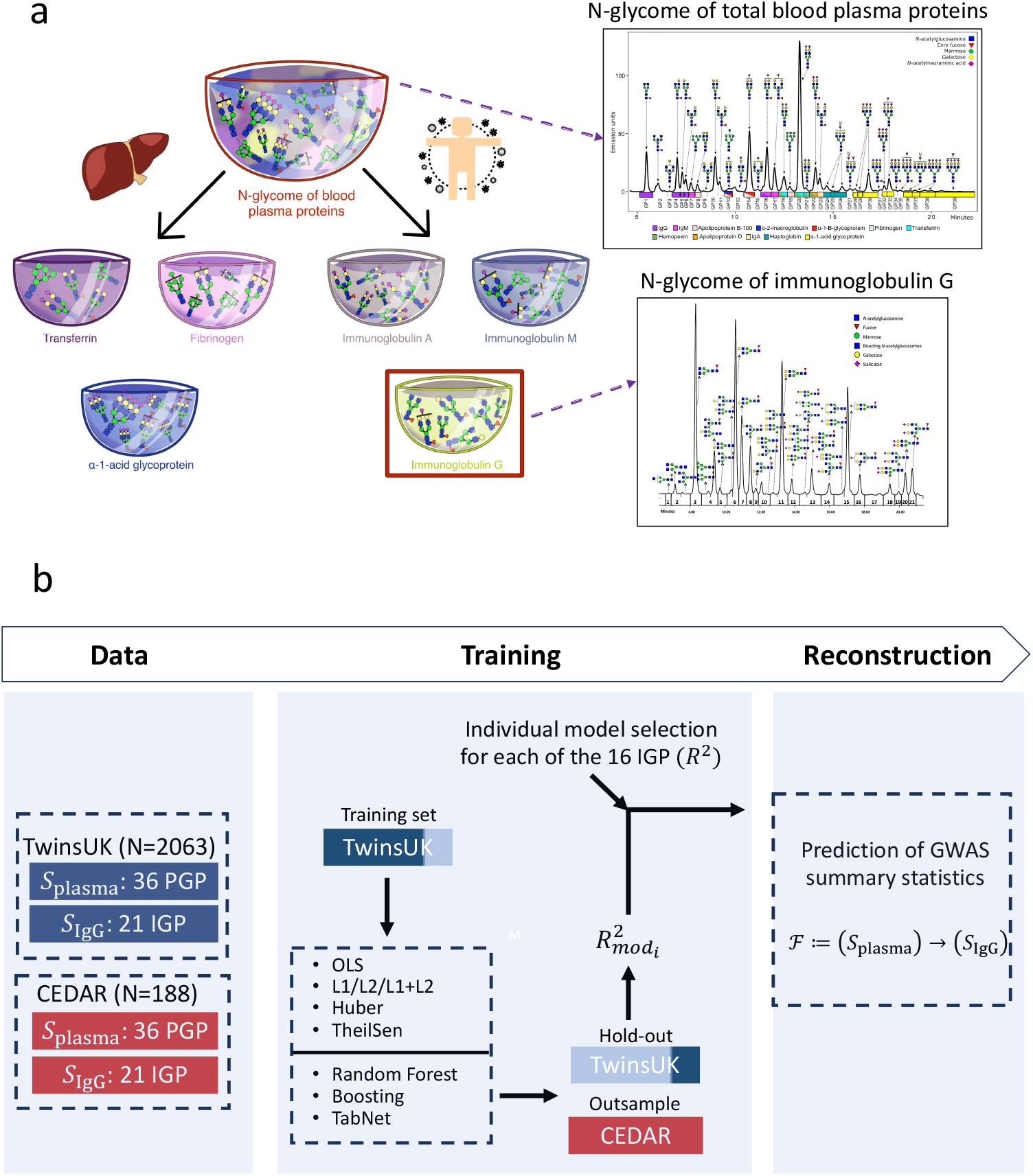
Prediction of IgG N-Glycome from TPNG. (a) Composition of the human blood plasma N-glycome. The plasma N-glycome is a mixture of N-glycomes from individual abundant glycoproteins. Major contributors shown include IgG, synthesized in lymphoid tissue, and transferrin, synthesized in the liver, illustrating the systemic origin of plasma glycans. A representative sample of IgG N-glycome is adopted from Fig.1^27^, a representative sample of plasma N-glycome is reused from Fig.1^66^ (b) A framework of reconstructing IgG N-glycome GWAS results using the TwinsUK cohort for training and the CEDAR cohort for external validation. We compared the performance of multiple linear (e.g., OLS, regularized, robust) and non-linear (Random Forest, Boosting, TabNet) models, and subsequently applied the trained model to summary statistics from the recently published TPNG GWAS^22^. For the GWAS summary statistic reconstruction, we used models for 16 out of the 21 IgG N-glycan peaks, selected based on a high proportion of variance explained (*R*^2^ > 0,5). S_plasma_ – the spectrum of TPNG, S_IgG_ – the spectrum of IgG N-glycome.

The IgG molecule consists of two main regions: the Fab region, responsible for antigen binding, and the Fc region, which interacts with immune cells and complement proteins. Both regions can undergo N-glycosylation, but the most important site is located in the Fc region at Asn297. Glycosylation enhances the stability of IgG, modulates its effector functions, and influences its half-life in circulation^11,12^. Study of IgG N-glycosylation provides insights into the etiology and pathophysiology of B-cell-mediated diseases, cardiometabolic diseases, and inflammatory conditions. Understanding the mechanisms underlying IgG glycosylation is crucial for unraveling the complex interplay between IgG modifications, cellular functions, and disease processes. Consequently, IgG glycans are emerging as promising therapeutic targets^13–15^ and biomarkers^16–18^, with significant potential for applications in precision medicine.

Quantitative genetics provides a unique opportunity to identify new regulators of protein N-glycosylation. Genome-wide association study (GWAS) is a widely used approach to map genetic loci involved in the control of multifactorial diseases and complex traits. To perform N-glycosylation GWASes, a large cohort of samples measured for both their N-glycomes and their genomes is required. Nowadays, the GWASes of total blood plasma^19–22^, immunoglobulin G^23–28^, transferrin^29^, and immunoglobulin A^30^ N-glycosylation identified several loci, but the number of analysed samples is still limited: 10,000 samples for total blood plasma N-glycosylation GWAS^22^; 11,000 samples for IgG N-glycosylation GWAS^24^; 21,000 for IgG galactosylation GWAS^28^; 2,000 for TF N-glycosylation GWAS^29^; and 2,000 for IgA N-glycosylation GWAS^30^.

N-glycome profiling typically involves multiple steps: extraction of glycoproteins, enzymatic release of glycans, purification, and subsequent analysis using techniques such as mass spectrometry or high-performance liquid chromatography. Each of these steps requires specialized reagents, equipment, and expertise, contributing to the overall expense. Solving the problem of reconstructing N-glycomes of individual glycoproteins from the total blood plasma can significantly extend the N-glycome datasets without profiling new samples.

In this study, we aimed to increase the sample size available for the genetic analysis of IgG N-glycosylation by reconstructing IgG-specific N-glycome traits from the total plasma N-glycome (TPNG) of already profiled samples. Our approach consisted of three main steps. First, we built a model for converting the 36 plasma N-glycome traits to 21 IgG N-glycome traits. We trained and evaluated multiple models using raw N-glycome measurements from both isolated IgG and total blood plasma in the TwinsUK (N=2,063) and CEDAR (N=188) cohorts. Second, we adopted the trained model to be directly applied to the summary statistics from the world’s largest TPNG GWAS to reconstruct summary statistics for each of IgG N-glycosylation trait. Finally, we identified new loci and replicated them in independent cohorts with directly measured IgG N-glycome profiles. Finally, we performed an *in-silico* gene prioritization for newly discovered loci using eight independent prioritization approaches to identify novel regulators of N-glycosylation.

## Results

### IgG N-glycome reconstruction

To develop a predictive model, we leveraged two independent cohorts, TwinsUK (N=2,063) and CEDAR (N=188), in which both the total plasma N-glycome (TPNG) and the isolated IgG N-glycome were measured for the same individuals. A preliminary analysis found significant linear correlations between total plasma and IgG N-glycomes across most of the N-glycome traits (Supplementary Figure 1). This supports our hypothesis that the IgG N-glycome represents a linearly extractable component of the total plasma N-glycome, which is a mixture of various N-glycomes of individual glycoproteins.

We trained and compared the performance of various regression models to predict the IgG N-glycome based on the total plasma N-glycome using 80% of the TwinsUK sample (*N* = 2063 * 0.8 = 1650). Each IgG N-glycan peak was predicted independently from the 36 total plasma N-glycan peaks (for a detailed description of the peaks, see Supplementary Table 1). We evaluated four regression models, including ordinary least squares (OLS), Elastic Net (with L1, L2, and combined L1/L2 regularization), Huber regression, and the Theil-Sen estimator. The performance of the models on the 20% hold-out TwinsUK set (*N* = 2063 * 0.2 = 413) is shown in Figure 2a. The coefficient of determination (R^2^) was used as the primary metric to assess the performance of the models. The results demonstrate substantial variation in prediction accuracy across the 21 IgG peaks – from R^2^ = 0,05 for IGP19 to R^2^ = 0,83 for IGP3 (Figure 2a).

**Figure 2:**
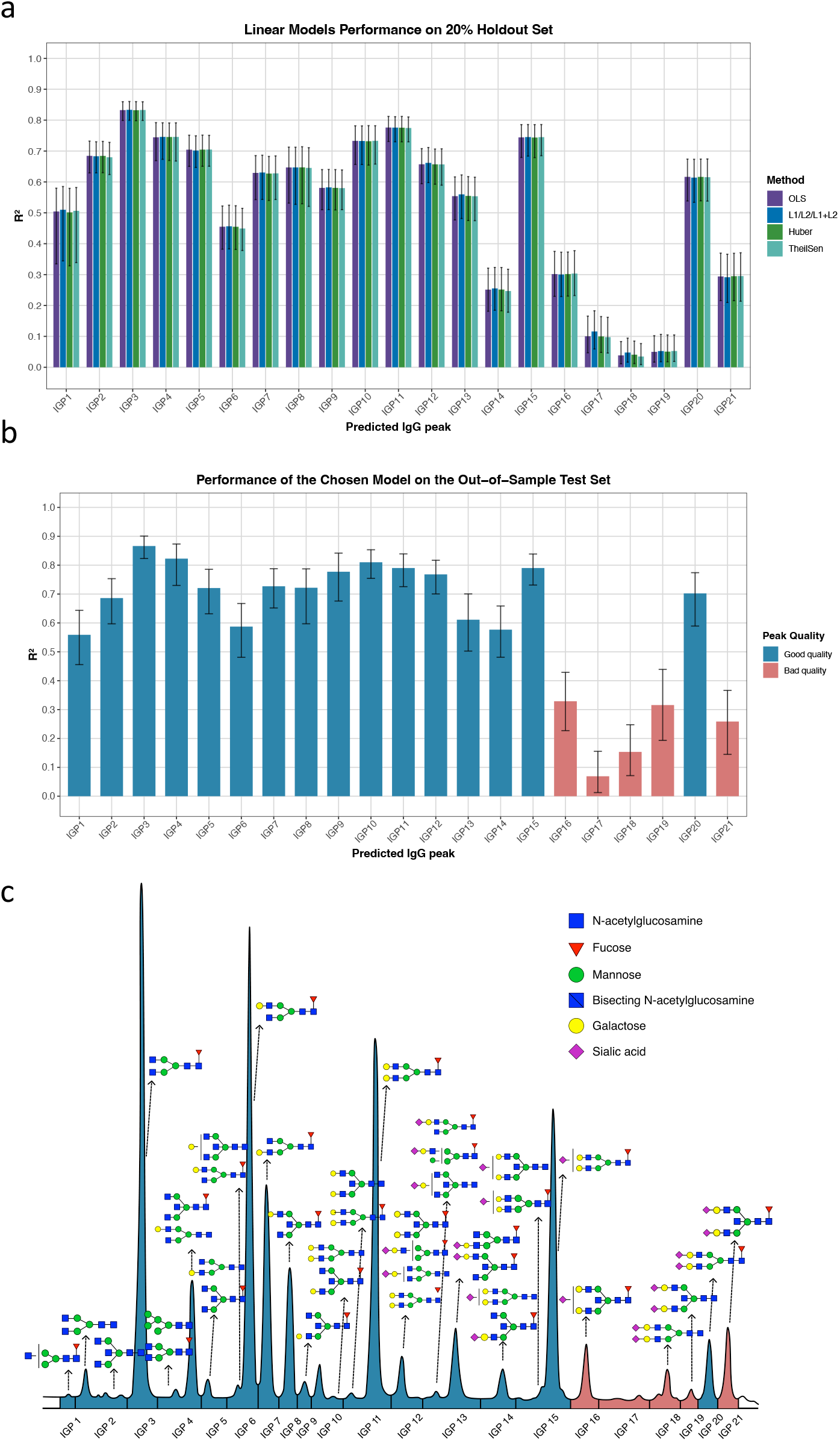
Comparative performance of linear models in predicting IgG glycan peaks from total plasma N-glycome data. Four models—ordinary least squares (OLS), elastic net framework (L1/L2/L1+L2), Huber regression and Theil-Sen Estimator —were evaluated. The plot shows (a) R^2^ values as evaluated on 20% hold-out from the TwinsUK cohort and (b) R^2^ values for the trained model on the hold-out, as calculated on the out-of-sample test set. (c) A representative IgG N-glycome profile depicting 16 glycan peaks with high reconstruction quality (blue) and 5 peaks with low reconstruction quality (red).

To test the sufficiency of the linearity assumption, we trained four non-linear models (namely, Random Forests (RF), Gradient Boosting (both Linear Regression Boosting and Tree Boosting), and the deep learning-based tabular model TabNet) and evaluated their performance on the 20% holdout from the TwinsUK. For each individual peak, the peak-specific linear model was selected and compared against the four non-linear models. Hereafter, the peak-specific linear model will be termed the trained model for simplicity. This comparison is presented in Supplementary Figure 2. More detailed description of the performance of linear and non-linear models are provided in Supplementary Table 2. Generally, non-linear predictors did not demonstrate an increase in performance, with the only exception of gradient boosting having marginally higher R^2^ for two peaks (IGP6 and IGP10) and random forest (IGP17 and IGP18).

We used the trained model on an independent test set (CEDAR cohort, N=188). IgG N-glycome prediction demonstrated robust performance on the test set (Figure 2b, Supplementary Table 2). To keep only traits with sufficient reconstruction accuracy for downstream analysis, we calculated a threshold by applying the Brent optimization method to identify distinct clusters of IgG N-glycome peaks with high and low accuracy. This approach identified a subset of 16 peaks (IGP1–IGP15 and IGP20), which formed a distinct cluster of IgG N-glycan peaks with high and reliable predictive performance. For the five peaks with low reconstruction accuracy (IGP16–IGP19 and IGP21), we may speculate that the particularly poor prediction is due to their origin from the Fab region of IgG rather than the Fc region. These Fab glycans likely constitute only a minor fraction of the corresponding total plasma N-glycome peaks, making their signal difficult to deconvolve from the more abundant glycoproteins in plasma.

In summary, the model performance on both the hold-out and independent test sets demonstrates that predicting the N-glycome of an individual glycoprotein, specifically IgG, is feasible for a significant part of the N-glycome spectrum of traits.

### Reconstructed IgG N-glycosylation GWAS

We reconstructed an IgG N-glycome GWAS to identify genetic variants specifically associated with IgG N-glycosylation. For this reconstruction, we applied the trained model to GWAS summary statistics from the recently published TPNG GWAS^22^. For the GWAS summary statistic reconstruction, we used models for 16 out of the 21 IgG N-glycan peaks, selected based on a high proportion of variance explained (*R*^2^ > 0.5) (Figure 2b).

The reconstructed GWAS identified 24 loci that passed the genome-wide significance threshold of *p=* 5 × 10^−8^/16 = 3.13 × 10^−9^, where 16 represents the number of N-glycome traits included in the analysis (Figure 3; Supplementary Data 3). Of these, 20 loci have been previously established as associated with IgG N-glycosylation^10^, confirming the high specificity and validity of our IgG N-glycosylation GWAS reconstructing approach.

**Figure 3:**
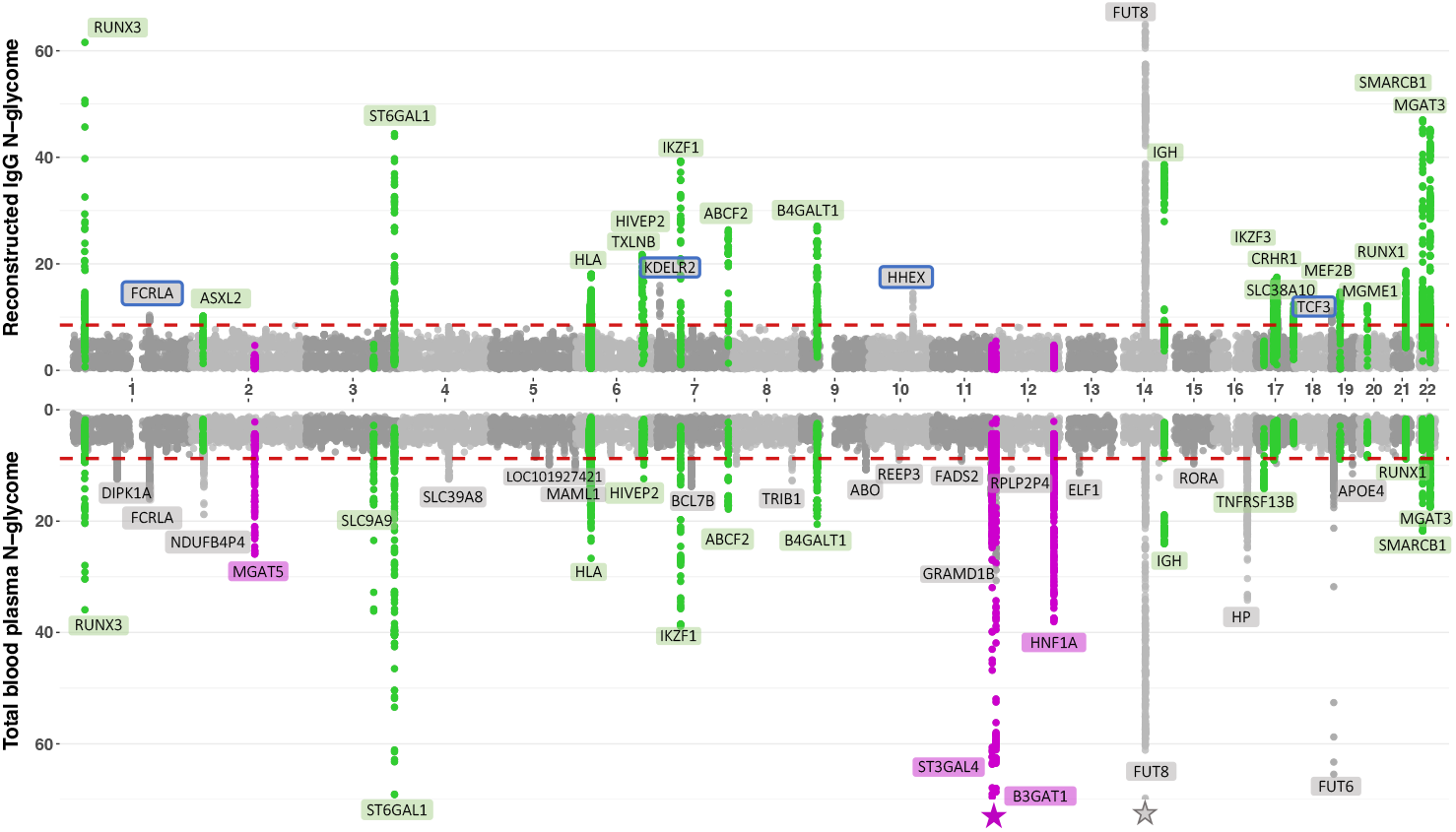
Miami plot of genome-wide association study results. The upper panel displays −log_10_(*P*) for associations from the reconstructed IgG N−glycome GWAS. The lower panel displays results for a total blood plasma N−glycome GWAS. Each point represents a single nucleotide polymorphism (SNP). Loci are alternated in color (transferrin-specific-purple; IgG-specific - green). The horizontal dashed lines indicate the genome-wide significance thresholds (5 × 10^−8^/16 for upper plot, 5 × 10^−8^/28 for bottom plot). Novel loci are highlighted with a box.

Interestingly, association signals for the majority of known IgG-specific loci were substantially more significantly associated with the reconstructed IgG N-glycosylation (16 N-glycome traits) compared to the source TPNG GWAS (N=10,764)^22^ and showed comparable significance to a genetic association signals from GWAS of 23 directly measured IgG N-glycome traits (N=11,237) from recently published studies^24,26^ (Figure 4, Supplementary Table 5). A prominent example is the *TXLNB* locus, whose association significance increased from *P* = 8.63 × 10^−9^ in the TPNG GWAS to *P* = 1.75 × 10^−22^ in the reconstructed IgG GWAS, demonstrating a marked gain in statistical power upon deconvolution of the plasma background. A similar pattern of enhanced significance was observed for 19 out of 28 loci containing IgG regulator genes (Supplementary Table 5).

**Figure 4:**
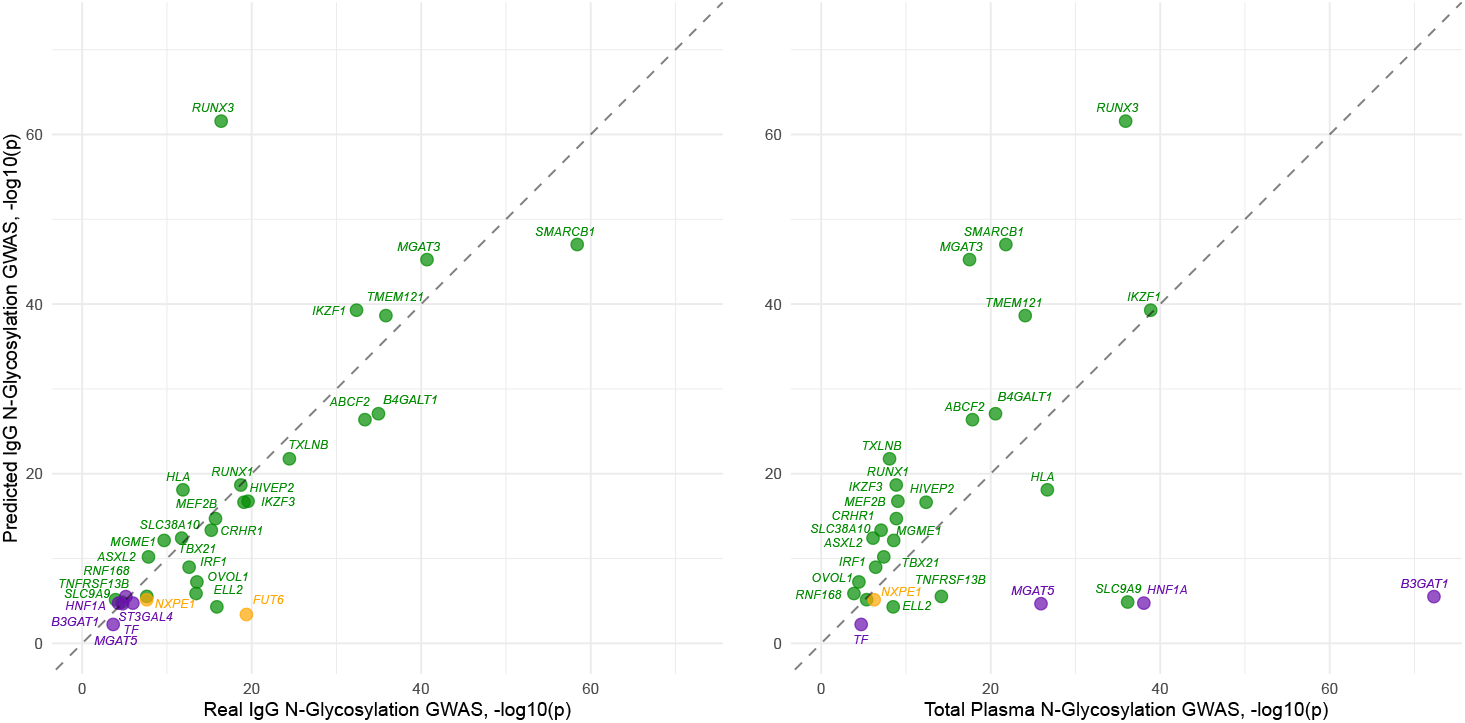
Comparative Analysis of P-Values from reconstructed IgG N-Glycosylation GWAS, real IgG N-Glycosylation GWAS, and source plasma N-Glycome GWAS. Data points represented as −log_10_(*P*) of three key GWASes correspond to known IgG/transferrin genetic loci, color-coded by source: IgG-associated (green), transferrin-associated (purple), and loci significant for both traits (orange).

Notably, loci specific to transferrin N-glycosylation regulation (*MGAT5, ST3GAL4, B3GAT1, HNF1A*), which showed strong association in the total blood plasma GWAS, were not associated with the reconstructed IgG N-glycome traits, serving as an effective negative control.

Four loci represent novel associations with the IgG N-glycome: *FCRLA, KDELR2, HHEX*, and *TCF3*. We successfully replicated two of these novel loci, *KDELR2* and *HHEX*, in an independent GWAMA of directly measured IgG N-glycome traits (see Methods), providing strong support for their role in the regulation of IgG N-glycosylation. The loci *FCRLA* and *TCF3* were not formally replicated, likely due to limited coverage in the replication cohort. While the maximum sample size in the replication cohort was 6,758, the sample size for the top SNPs at these loci was only ∼2,500, yielding an estimated replication power of approximately 60%. For the *TCF3* locus, when considering only SNPs with sufficient coverage in the replication dataset, the lead SNP became rs4807942 (discovery *P* = 5.69 × 10^−11^, replication *P* = 1.61 × 10^−7^). For the *FCRLA* locus, no genome-wide significant SNP from the discovery analysis was well-covered in the replication dataset. However, this locus was replicated in TPNG GWAS by Timoshchuk et al^31^.

Despite these convincing and largely replicable GWAS results, we noted a strong variability in the genomic control inflation factor (λ), ranging from 0.72 for IGP4 to 1.15 for IGP13 (Supplementary Figure 3, Supplementary Table 4). This contrasts with conservative λ values (range: 1.01 to 1.05) observed in the original TPNG GWAS summary statistics used as input (Supplementary Data 4). QQ-plots for each glycan peak revealed deflections below the diagonal for peaks with particularly low λ values. We posit that this pattern of inflation/deflation of genetic association test statistics is an artifact of the summary statistic reconstruction process itself and is not directly indicative of violation of genetic association analysis assumptions, e.g., population stratification or polygenicity. To further evaluate the consistency of the reconstructed effects, we compared effect sizes of genetic variants on N-glycome traits (beta-beta plot, Supplementary Figure 4) between the discovery (reconstructed IgG GWAS) and replication (directly measured IgG GWAS) for SNPs that reached genome-wide significance in either analysis. The slope of the regression line varied from 0.528 to 3.083 across the 16 peaks, where a slope of 1 would indicate perfect concordance.

In summary, our reconstructed IgG N-glycosylation GWAS produced clear and biologically meaningful results. The approach successfully recovered known IgG glycosylation loci with stronger association signals than those observed in the TPNG GWAS, while effectively filtering out associations attributable to N-glycome traits of other liver-derived plasma proteins. Furthermore, we identified four novel genetic loci: *FCRLA, KDELR2, HHEX*, and *TCF3*. Notably, performing the GWAS using only 16 direct reconstructed IgG peaks was sufficient to detect associations with most established IgG glycosylation loci (20 out of 29^10^). This demonstrates greater analytical efficiency compared to conventional study designs, which typically require analysis of more direct glycan peaks (e.g., 23) and numerous derived traits (up to 54).

### Gene prioritization for newly discovered loci

Identification of genes, rather than genetic loci, can help to find novel protein glycosylation regulators and suggest targets for intervention in glycome-related diseases. We prioritized candidate genes using a consensus approach integrating eight sources of evidence - based on a literature search of genes encoding known enzymes and regulators of N-glycan biosynthesis; genes causing congenital disorders of glycosylation; colocalization of glyQTLs with eQTLs and blood plasma pQTLs; annotation of putative causal variants affecting protein structure; enrichment of gene sets and tissue-specific expression; and prioritization of the nearest gene (see Methods).

We prioritized three candidate genes in four glyQTLs (Supplementary Data 6a) by selecting the gene with the highest unweighted sum of evidence across all eight predictors^32^: *KDELR2, HHEX*, and *TCF3*. Two of these genes, *HHEX* and *TCF3*, encode transcription factors involved in lymphopoiesis^33–37^. *TCF3*, commonly known as E2A, encodes a basic Helix-Loop-Helix transcription factor that functions as E12/E47 homo- or heterodimers, with both monomers originating from alternative splicing of a single gene. E2A is a master-regulator of lymphoid lineage commitment, controlling the B-cell maturation and T-cell progenitor differentiation. Insufficiency of either isoform hinders progression through B-cell development stages^33–35^. Additionally, E2A regulates activity of *RAG1* and *RAG2* genes, which mediate V(D)J recombination in immunoglobulin and TCR genes, deficiency of which causes immunological disorders^33,36,37^. *HHEX* (Hematopoietically Expressed Homeobox) encodes a transcription factor expressed in early lymphoid progenitors, with expression downregulated upon terminal differentiation^38,39^. In murine models, *HHEX* depletion results B-cell developmental defects beginning at the pro-B-cell stage, and impairs T- and B-cells repopulation^40^. Human evidence remains limited, although *HHEX* is among the lead gene signature of T-cell acute lymphoblastic leukemia^41^. Beyond lymphopoiesis, *HHEX* functions in early embryonic development and organogenesis of liver, thyroid, and lung^42^, suggesting potential systemic contributions to glycosylation regulation. These findings indicate that these loci may influence IgG N-glycosylation through cell differentiation state control, although direct control of glycan biosynthesis genes cannot be ruled out. *HHEX* and *TCF3* extends the list of established lymphoid transcription factors implicated in IgG N-glycosylation, which already includes *IKZF1* and *IKZF3*, and *RUNX1* and *RUNX3*^24^.

Another prioritized gene, *KDELR2*, encodes a KDEL receptor mediating retrograde transport of ER lumen resident proteins from the Golgi. The KDEL-retrieval system maintains ER localization of chaperons and enzymes critical for N-glycan processing and glycoprotein quality control, including BiP, calreticulin, and UGGT^43,44^. Genetic variation affecting this system is thus directly relevant to glycosylation machinery retention, and consequently N-glycosylation levels.

Locus label *FCRLA* yielded no genes we could prioritize in the study. However, this locus has been previously identified and replicated in total plasma N-glycosylation GWAS^31^. The region is enriched for Fc-receptors genes, including the classic Fc-gamma receptor (*FCGR1A, FCGR2A, FCGR2B, FCGR3A, FCGR3B*), the ligand-binding alpha chain of the high-affinity IgE receptor (*FCER1A*), the Fc-receptor-like genes (*FCRL1-5*), and *FCER1G*, the gene encoding the gamma chain of the Fc-epsilon receptor, a signaling component shared by Fc-epsilon-RI activating Fc-gamma receptors^45^. Several Fc-gamma receptor genes showed eQTL colocalization with glyQTLs (Supplementary Figure 5A), although these eQTLs derive from whole blood or non-B-cell populations (granulocytes, T cells) rather than B cells or plasma cells. The locus also harbours *B4GALT3*, encoding a beta-1,4-galactosyltransferase directly involved in glycan biosynthesis, and *FCRLA*, which had the highest polygenic priority score (PoPS, see Methods). *FCRLA* encodes a soluble ER-retained receptor capable of immunoglobulin binding with hypothesized roles in both germinal center B-cell differentiation and direct modulation of plasma N-glycosylation through ER retention of immunoglobulins, predominantly IgM^46,47^. A previous study prioritized *FCGR2B* as the most likely effector gene, focusing on its role in inhibiting B-cell activation, which could decrease antibody production and thus influence N-glycome^48,49^. The absence of a clearly prioritized gene suggests a more complex architecture: the convergence of Fc receptors, a galactosyltransferase, and a B-cell differentiation factor within a single associated locus suggests multiple potential causal mechanisms, which the present study could not resolve.

Analysis of the PoPS model features most informative in prioritizing IgG N-glycosylation-relevant genes yielded enrichment for pathways plausibly influencing N-glycosylation. The highest-ranking cluster contained features related to B- and T-cell differentiation and developmental transcription factors, concordant with the prioritization of *TCF3* and *HHEX*, and the hypothesized role of *FCRLA* in B-cell differentiation into memory and plasma cells^50^. Other top clusters involved Rho-GTPase-mediated immune receptor signalling, transcriptional regulation and chromatin remodelling, and MAPK-mediated growth-factor signalling (Supplementary Figure 5E) – pathways, relevant to immune cell activation and Fc receptor function, consistent with the *FCRLA* locus content. Together, these findings further support the coherence of prioritized genes, linking IgG N-glycome variation to lymphocyte differentiation and immune signalling.

## Discussion

Harnessing the genetic regulation of protein-specific glycosylation has been constrained by the practical limitations of large-scale, targeted glycomics. Here, we present a computational framework that transcends this barrier by reconstructing the IgG N-glycome from the total plasma N-glycome. Our trained model accurately reconstructs IgG-specific glycosylation traits and when applied to GWAS summary statistics, unveils genetic regulators with enhanced specificity and power. This method allows conducting research of IgG N-glycome by leveraging existing TPNG data, enabling discoveries without the need to profile novel samples.

Our comparative analysis of linear and non-linear models revealed that linear models are sufficient for high-fidelity reconstruction of most Fc-region IgG glycans from the TPNG. This finding strongly supports the hypothesis that IgG glycosylation contribution to the plasma glycome mixture is a linearly separable component. The failure to accurately reconstruct a subset of peaks (IGP16–IGP19, IGP21) is itself informative, pointing to their likely origin from the Fab region, where they represent a minor fraction of the corresponding total plasma signal.

The most compelling validation of our framework comes from the reconstructed IgG N-glycosylation GWAS. The recovery of 20 out of 24 significant loci as known IgG regulators demonstrates remarkable specificity. Crucially, the association signals for the most loci (Supplementary Data 5) like *TXLNB, RUNX3*, and *ABCG2* were markedly enhanced in our reconstructed GWAS compared to the source TPNG GWAS, and reached significance levels comparable to direct IgG GWAS. This “signal purification” effect—whereby the IgG-specific genetic signal is amplified by removing the noise and confounding contributions from other plasma glycoproteins—is a central showcase of our method. Conversely, the absence of transferrin-specific loci (*MGAT5, ST3GAL4*) serves as a powerful negative control, confirming that our reconstruction is not a mere recapitulation of the TPNG GWAS but a targeted extraction of its IgG part.

We identified and prioritized four novel candidate regulators of IgG glycosylation: *FCRLA, KDELR2, HHEX*, and *TCF3*. The biological plausibility of these candidates strengthens our findings: *FCRLA* is an endoplasmic reticulum-resident protein expressed in B-cells, intimately involved in IgG assembly; *KDELR2* is a retrograde transport receptor crucial for Golgi-ER trafficking of glycosylation enzymes; *HHEX* and *TCF3* are transcription factors with established roles in B-cell development and function. This suggests that genetic variation influences IgG glycosylation not only through canonical glycosyltransferases but also via regulators of B-cell biology and intracellular protein trafficking, opening new mechanistic avenues for exploration.

Because reconstructed effects are derived through peak-specific linear models, effect-size magnitudes are not guaranteed to be on the same scale as directly measured IgG GWAS; we therefore interpret reconstructed GWAS primarily for locus discovery and effect direction. Indeed, the variable concordance of effect sizes of found loci on reconstructed IgG N-glycome traits and directly measured N-glycome traits highlights that our method optimizes for the accurate detection and direction of associations, while the precise estimation of effect magnitudes may be influenced by the reconstruction model’s coefficients. Future work incorporating individual-level genotype data could refine these estimates.

The implications of this study are threefold. First, for the field of glyco-genetics, we provide a readily applicable tool to generate protein-specific glycome GWASs from existing TPNG datasets, substantially expanding the discovery potential. Second, for functional genomics, we delivered new high-confidence candidate genes (*FCRLA, KDELR2, HHEX, TCF3*) for experimental follow-up to dissect their precise roles in the IgG glycosylation pathway. Third, for translational research, enhanced understanding of IgG glycan regulation offers new levers for therapeutic intervention in antibody-mediated diseases and improves the utility of IgG glycosylation as a biomarker by clarifying its genetic determinants.

In conclusion, we proposed a method that turns the complexity of the plasma glycome— a mixture of glycans from various glycoproteins—from a challenge into an opportunity. By computationally deconvoluting it, we can now leverage the vast investment in TPNG studies to power the genetic discovery of protein-specific glycosylation, bringing us closer to a functional understanding of this critical post-translational modification and its role in human health and disease.

## Materials and methods

### Study datasets description

For the IgG N-glycome reconstruction from the total blood plasma N-glycome we used TwinsUK and CEDAR datasets with both measured TPNG and IgG glycosylation profiles.

#### TwinsUK

The TwinsUK cohort^51^, also referred to as the UK Adult Twin Register, is a nationwide registry of volunteer twins in the United Kingdom. The Department of Twin Research and Genetic Epidemiology at King’s College London (KCL) hosts the registry. From this registry, 2,063 subjects had N-linked total plasma glycan and immunoglobulin G measurements, which were included in the analysis. The analysis included 1953 women and 110 men, averaging 52 years of age (range: 19-83).

#### CEDAR

The Correlated Expression and Disease Association Research (CEDAR) study was described previously^52^. Peripheral blood as well as intestinal biopsies (ileum, transverse colon, rectum) were collected from 188 healthy Europeans visiting the Academic Hospital of the University of Liège as part of a national screening campaign for colon cancer. Participants included 108 women and 80 men, averaging 55 years of age (range: 18-85). Enrolled individuals were not suffering any autoimmune or inflammatory disease and were not taking corticosteroids or non-steroidal anti-inflammatory drugs (except for low doses of aspirin to prevent thrombosis).

### Plasma N-glycome quantification

Plasma N-glycome quantification of both TwinsUK and CEDAR samples was performed at GENOS by applying the following protocol. Plasma N-glycans were enzymatically released from proteins by PNGase F, fluorescently labeled with 2-aminobenzamide, and cleaned up from the excess of reagents by hydrophilic interaction liquid chromatography solid phase extraction (HILIC-SPE) as described previously^53^. Fluorescently labeled and purified N-glycans were separated by HILIC on a Waters BEH Glycan chromatography column, 150 × 2.1 mm, 1.7 μm BEH particles, installed on an Acquity ultra-high-performance liquid chromatography (UHPLC) instrument (Waters, Milford, MA, USA) consisting of a quaternary solvent manager, sample manager, and a fluorescence detector set with excitation and emission wavelengths of 250 nm and 428 nm, respectively.

### IgG N-glycome quantification

#### TwinsUK sample

Isolation of IgG and glycan analysis has been previously described in^24^. IgG was isolated from human plasma using protein G monolithic plates. Following elution with formic acid and neutralization, N-glycans were released from the purified IgG by incubation with PNGase F after protein denaturation. The released glycans were fluorescently labeled with 2-aminobenzamide (2-AB) and purified via hydrophilic interaction solid-phase extraction (HILIC-SPE) to remove excess reagents. The labeled N-glycans were separated by HILIC-UPLC with fluorescence detection, and chromatograms were processed to quantify 24 individual glycan peaks. From these primary peaks, 21 glycosylation traits were calculated based on a unified set of peaks that were matched between the CEDAR and TwinsUK samples (Supplementary Table 1a).

#### CEDAR sample

IgG N-glycome quantification of the CEDAR samples was performed at Genos by applying the following protocol. IgG N-glycans were enzymatically released from proteins by incubation with PNGase F, fluorescently labelled with 2-aminobenzamide, and cleaned up from the excess of reagents by hydrophilic interaction liquid chromatography–solid-phase extraction (HILIC–SPE). The fluorescently labelled and purified N-glycans were separated by HILIC on a Waters BEH Glycan chromatography column, 150 × 2.1 mm, 1.7 μm BEH particles, installed on an Acquity UPLC instrument (Waters, Milford, MA, United States) consisting of a quaternary solvent manager, a sample manager, and a fluorescence detector set with excitation and emission wavelengths of 250 and 428 nm, respectively. IgG N-glycans were separated into 22 peaks. From these primary peaks, 21 glycosylation traits were calculated based on a unified set of peaks that were matched between the CEDAR and TwinsUK samples (Supplementary Table 1a).

### Harmonization and normalization of N-glycan peaks

To prepare training and test datasets for IgG N-glycome reconstruction, we harmonized the set of glycan peaks by applying a protocol published in^20^. This same protocol was also used in the work by Sharapov et al^22^, from which we obtained the GWAS results for total blood plasma N-glycosylation (summary statistics) for the second part of our study. We applied a harmonization procedure that resulted in a set of 36 plasma GPs (PGP) and 21 IgG GPs (IGP) for both datasets (see Supplementary Table 1 a, b).

Normalization and batch correction were performed on harmonized UHPLC glycan data separately for both cohorts using the protocol published in^21^. We used probabilistic median quotient normalization^54,55^. This approach calculates the dilution factor of each sample relative to a reference sample, defined as the median abundance of each glycan peak across all samples. For each sample, a vector of quotients is created by dividing each GP measurement by the reference value. The median of these quotients serves as the dilution factor, and original sample values are divided by it. The assumption is that varying intensities among individuals correspond to different amounts of biological material in the samples.

Normalized glycan measurements were log-transformed due to the right skewness of distribution and multiplicative nature of batch effects. Outlying measurements were removed. Outlier was defined as a sample that had at least one GP that was greater than upper quartile + 3 * IQR (interquartile range) or lower than lower quartile - 3 * IQR. Finally, sex and age adjustments were made, and the data were standardized to a mean of 0 and a standard deviation of 1.

### Computational reconstruction of N-glycome: formulation and methods

The problem of reconstructing the 21 IgG N-glycan traits from the 36 total plasma N-glycan traits can be seen as 21 independent regression problems. Let PGP1-PGP36 represent the normalized concentrations of the 36 glycan peaks derived from the total blood plasma N-glycome. The objective was to train models that, given these 36 plasma peaks, would accurately predict the values of IGP1-IGP21 corresponding to 21 glycan peaks from the isolated IgG N-glycome.

Given the anticipated linear relationships between the plasma and IgG glycan abundances, our primary focus was on linear models. We trained and evaluated the performance of four linear regression models: (a) Ordinary Least Squares (OLS, serving as baseline); (b) elastic-net regularized regression, spanning the full mixing range to include the L1-only (Lasso) and L2-only (Ridge) limits; (c) L2-regularized Huber regression; and (d) Theil-Sen robust regression (multivariate generalization of the Theil– Sen estimator). To assess the robustness of our prespecified linearity assumption, we fitted four non-linear models: Random Forests (RF), Gradient Boosted Tree and Linear Regression, and a deep learning-based tabular model, TabNet.

Hyperparameters were tuned via five-fold cross-validation on an 80% training subset of the TwinsUK cohort (80% out of N=2063). Grid search was used for linear models, randomized search for RF, and Bayesian optimization with the Tree-structured Parzen Estimator sampler via the optuna library^56^ for the boosting models and TabNet. Subsequently, models were refit on the full 80% training split using the selected optimal hyperparameters. The best-performing model was selected based on the 20% holdout performance based on the greatest *R*^2^. Final generalization performance was assessed on the independent CEDAR cohort (N=188) with no further tuning. Non-linear models were treated strictly as sensitivity analyses, and we report only their performance on the holdout.

### Genetic association analysis

#### Discovery

To identify genetic loci associated with IgG N-glycosylation, we reconstructed GWAS summary statistics of 16 selected IgG N-glycan traits with high predictive accuracy (*R*^2^ > 0.5 on the CEDAR cohort).

The reconstruction of GWAS summary statistics for the IgG N-glycan traits was performed by applying the pre-trained prediction models (described in the previous section) directly to the summary-level statistics from the largest available GWAS of total blood plasma N-glycosylation^22^. Specifically, we used the effect estimates (betas) and standard errors for the 36 total plasma glycan peaks (PGP1–PGP36) from the combined discovery and replication meta-analysis of 10,764 individuals (as detailed in the original study). The genetic association signals for the reconstructed IgG glycans were obtained using linear transformation described in Shadrina et al^26^:

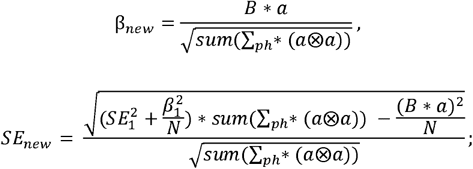

where *B*— matrix, where each column *β*_*i*_ — effect of each SNP in GWAS_i_; *α* — vector of linear combination coefficients; ∑_*ph*_ — matrix of correlations between 36 plasma N-glycosylation traits; ⊗ — outer product; *SE*_1_ — standard error of each SNP’s effect in the first GWAS_1_; *N* — minimal N out of all GWASes for each SNP.

These formulas allow for the calculation of GWAS summary statistics for a linear combination of traits. This approach allows us to avoid performing a GWAS on the reconstructed trait values using individual-level glycome and genetic data under the assumption of a linear relationship.

The sample characteristics, genotyping, imputation, and quality control procedures for the underlying plasma N-glycome GWAS are fully described in the original publication^22^. Briefly, the analysis included participants of seven cohorts (TwinsUK, EPIC-Potsdam, PainOmics, SOCCS, SABRE, QMDlab, and CEDAR), comprising a total sample size of 10,764 individuals, of whom 10,172 (94,5%) were of European ancestry and 592 (5,5%) were of non-European ancestry. Genotypes were imputed using the Haplotype Reference Consortium or 1000 Genomes Project panels. The meta-analysis was conducted using a fixed-effects inverse-variance weighted model.

In the original plasma N-glycome GWAS, the genomic control inflation factor (λ) for the 36 traits ranged from 1.01 to 1.05, indicating minimal population stratification. In our analysis of the 16 reconstructed IgG N-glycan traits, the genomic control inflation factor ranged from 0.72 to 1.15 (Supplementary Figure 3). We contend that this inflation/deflation of λ is a methodological artifact of the reconstruction procedure itself and does not provide evidence for true population stratification or conventional polygenicity.

To define genome-wide significant glyQTLs, we used a conventional genome-wide significance threshold, Bonferroni corrected by 16 IgG N-glycan traits (*P* ≤ 3.13 × 10^−9^). Loci were defined as genomic regions within a 250 kb window of the lead single-nucleotide polymorphism (SNP).

#### Replication

To replicate the genetic associations identified in the discovery analysis, we performed a replication GWAS using independent summary statistics derived from a set of cohorts from Klarić et al^24^ and Shadrina et al^26^ studies. The replication summary statistics were generated by meta-analyzing GWAS results from seven independent cohorts: CROATIA-Korčula (N = 849), CROATIA-Vis (N = 802), ORCADES (Orkney Complex Disease Study) (N = 1960), EGCUT (N=575), FINRISK (N=552), CROATIA-Korčula2 (N=941) and VIKING (N=1079). The total sample size of the replication cohort was 6,758 individuals, all of European ancestry. The sample characteristics, genotyping, imputation, and quality control procedures for these cohorts are fully described in the original studies (Klarić et al. for CROATIA-Korčula, CROATIA-Vis and ORCADES cohorts and Shadrina et al. for EGCUT, FINRISK and CROATIA-Korčula2 cohorts).

The original analyses were based on 23 directly measured IgG N-glycosylation traits. To ensure consistency with our discovery analysis, we mapped the summary statistics from the original 23 traits to the set of 16 selected IgG N-glycan peaks used in our study, based on the mapping provided in Supplementary Data 1. A fixed-effects, inverse-variance-weighted meta-analysis was conducted to combine the GWAS summary statistics across the six replication cohorts. The genomic control inflation factor (λ) for the 16 replicated traits in this meta-analysis ranged from 0,81 to 1,00.

Loci were considered successfully replicated if the lead SNP showed a statistically significant association with the same IgG N-glycan trait in the replication meta-analysis at a Bonferroni-corrected threshold of *P* < 0,0125 (0,05/4, where 4 is the number of discovered novel loci). The direction of the effect (beta) for the effect allele was consistent between the discovery and replication analyses. The results of the replication analysis, including lead SNP information, effect sizes, and P-values, are presented in Supplementary Table 3.

### Prioritization of candidate genes in found loci

For the purpose of post-GWAS analyses, we used reconstructed discovery summary statistics (N=10,172). We applied an ensemble of methods to prioritize plausible candidate genes in the four discovered loci. We applied eight approaches to prioritize the most likely effector genes: (1) prioritization of the nearest gene; (2) prioritization of genes with known role in biosynthesis of N-glycans; (3) genes of congenital disorders of glycosylation; (4) genes with direct experimental support for regulation of protein N-glycosylation; (5) prioritization of genes containing variants in strong LD (*R*^2^ ≥ 0.8) with the lead variant, which are protein truncating variants (annotated by Variant Effect Predictor, VEP^57^) or predicted to be damaging by FATHMM XF^58^, FATHMM InDel^59^; (6) prioritization of genes whose eQTL and/or (7) pQTL are colocalized with glyQTL; (8) prioritization of genes based on the PoPS framework^60^. We prioritized the most likely ‘causal gene’ for each association using a consensus-based approach, selecting the gene with the highest, unweighted sum of evidence across all eight predictors. In the case of equality of the scores for two genes, we prioritized both genes, but we selected the random one to represent the locus in the tables/figures to avoid cluttering.

#### Functional annotation of genetic variants

We inferred the possible molecular consequences of genetic variants in glyQTLs. We focused on variants in LD with lead (for univariate and multivariate signals) and sentinel variants (for univariate signals) picked by COJO. We created a “long list” of putative causal variants using PLINK version 1.9 (--show-tags option), applied to whole genome re-sequenced data for 503 European ancestry individuals (1000 Genomes phase 3 version 5 data). The size of the window to find the LD in both cases was equal to 500kb. The default value was taken as a threshold to include SNPs into the credible sets. Ensembl Variant Effect Predictor (VEP) (Supplementary Data 6d) and by FATHMM-XF (Supplementary Data 6b), FATHMM-INDEL (Supplementary Data 6c) to reveal pathogenic point mutations.

#### Genes of N-glycan biosynthesis and Congenital Disorders of Glycosylation

We searched for the genes encoding glycosyltransferases—enzymes with a known role in N-glycan biosynthesis^61^—located in the ±250 Kb vicinity of the lead SNPs in glyQTLs. Additionally, we prioritized genes with known mutations that cause Congenital Disorder of Glycosylation according to the MedGen database (https://www.ncbi.nlm.nih.gov/medgen/76469) that are located in the vicinity of ±250kb from the lead SNPs.

#### Colocalization with eQTL and pQTL

To identify potential pleiotropic effects of glyQTL on gene expression in relevant tissues, we applied Summary data-based Mendelian Randomization (SMR) analysis followed by the Heterogeneity in Dependent Instruments (HEIDI) test^62^. Expression QTLs (eQTLs) were obtained from three collections: Westra et al^63^ (peripheral blood), GTEx v7^64^ (whole blood), CEDAR^52^ (CD19+ B lymphocytes, CD8+ T lymphocytes, CD4+ T lymphocytes, CD14+ monocytes, CD15+ granulocytes); and a collection derived from antibody-based Olink Explore assay^65^ for protein QTLs (pQTLs). The outcome variable was the reconstructed IgG N-glycome trait with the strongest association. Only cis-QTLs, defined as associations within 1 Mbp of the nearest gene boundary, were considered for gene prioritization.

SMR results were considered statistically significant if P_adj < 0.05 (Benjamini-Hochberg adjusted p-value). For the HEIDI test, a P<0.05 threshold was used to reject the null hypothesis of no difference between association profiles (Supplementary Data 6e, 6f).

#### PoPS

Polygenic priority score (PoPS)^60^ was used for gene-set-based prioritization. Within each locus, the gene with the highest PoPS score was selected. To analyse feature contribution, 50 PCs were derived from the scaled gene by feature matrix using truncated SVD. Complete linkage hierarchical clustering, using squared Pearson correlation between PCs as a similarity measure, grouped features into clusters with *R*^2^ > 0.01. This threshold follows that of original authors^60^. See Supplementary Data 6g for details on cluster composition.

## Acknowledgments and Funding

The work of S.Sh., A.T., D.M., A.S., Y.S.A. was supported by the Research Program at the Moscow State University (MSU) Institute for Artificial Intelligence. The work of A.S. was also supported by Non-commercial Foundation for the advancement of Science and Education “INTELLECT”. C.M. is funded by the Chronic Disease Research Foundation (CDRF), by the Italian Ministry of Education and Research: Dipartimenti di Eccellenza Program 2023 to 2027 and by the Italian Ministry of Health – Bando Ricerca Corrente.

TwinsUK was funded by the Wellcome Trust, Medical Research Council, Versus Arthritis, European Union Horizon 2020, Chronic Disease Research Foundation (CDRF), Wellcome Leap Dynamic Resilience Programme (co-funded by Temasek Trust), Zoe Ltd, the National Institute for Health and Care Research (NIHR) Clinical Research Network (CRN) and Biomedical Research Centre based at Guy’s and St Thomas’ NHS Foundation Trust in partnership with King’s College London.

## Author contribution

A.S. coordinated this study; A.S., D.M., A.T., S.Sh. contributed to the design of the study, carried out statistical analysis; A.S., D.M., A.T. produced the figures; A.S., D.M., A.T., S.Sh., Y.S.A. contributed to interpretation of the results; A.S., D.M., A.T., S.Sh. wrote the first version of the manuscript; C.J.S. and C.M. contributed to interpretation of the results; M.G. designed CEDAR study and contributed to interpretation of the results; Y.S.A. and G.L. conceived and oversaw the study, contributed to the design and interpretation of the results; all co-authors contributed to the final manuscript revision.

## Abbreviations

IgG: immunoglobulin G
TPNG: Total Plasma N-glycome
IGP: Immunoglobulin G N-glycan peak
PGP: Total Blood Plasma N-glycan peak
GWAS: Genome-Wide Association Study

