## Supplementary Figures for "Reconstructing N-Glycan Profiles of Individual Glycoproteins from the Total Blood Plasma N-Glycome: Implications for Immunoglobulin G N-Glycosylation GWAS"

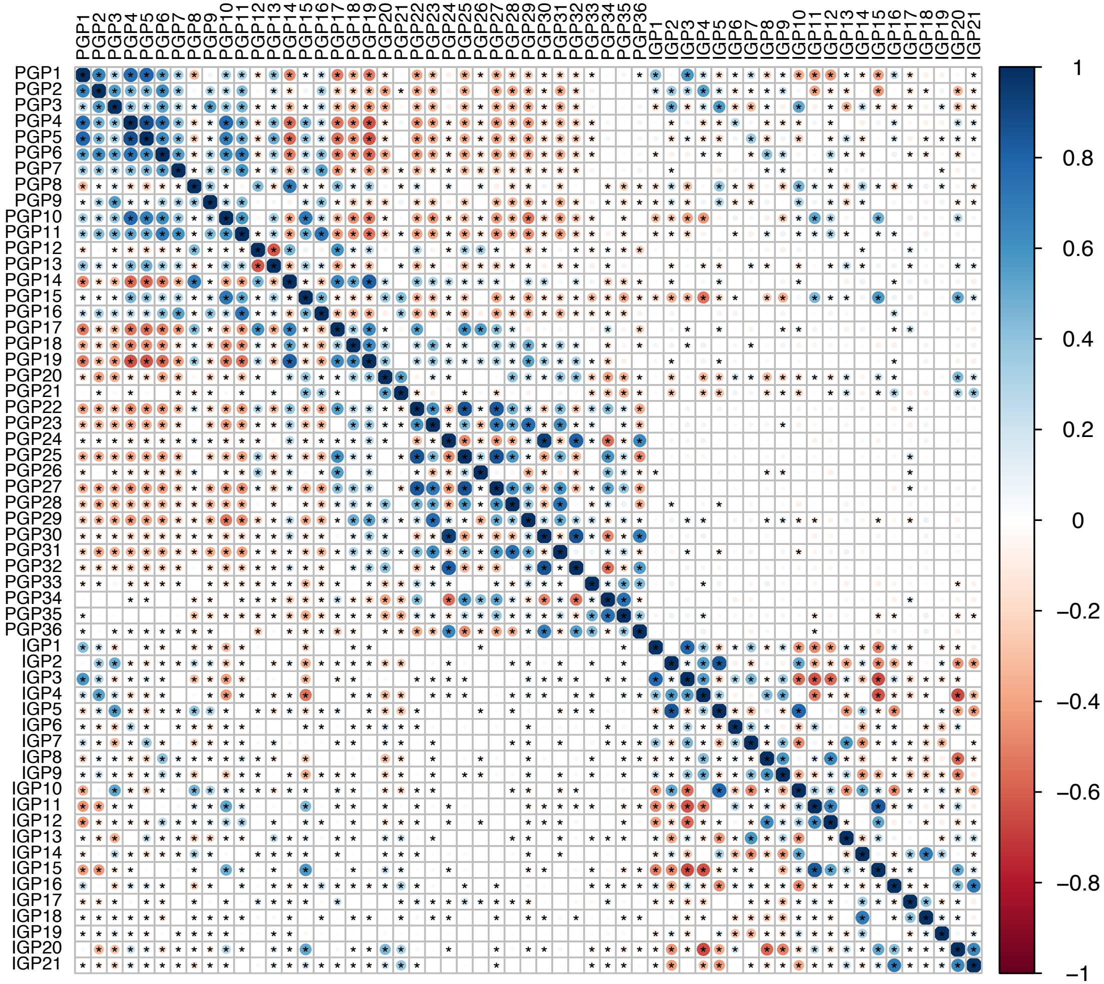

Supplementary Figure 1. Pearson correlation heatmap between plasma and IgG N-glycan peaks in the TwinsUK cohort. The heatmap depicts correlations across 36 total blood plasma N-glycan peaks and 21 IgG N-glycan peaks; cohort sample size N = 2063. Asterisks (*) indicate statistically significant correlations ($p < 0.05$).

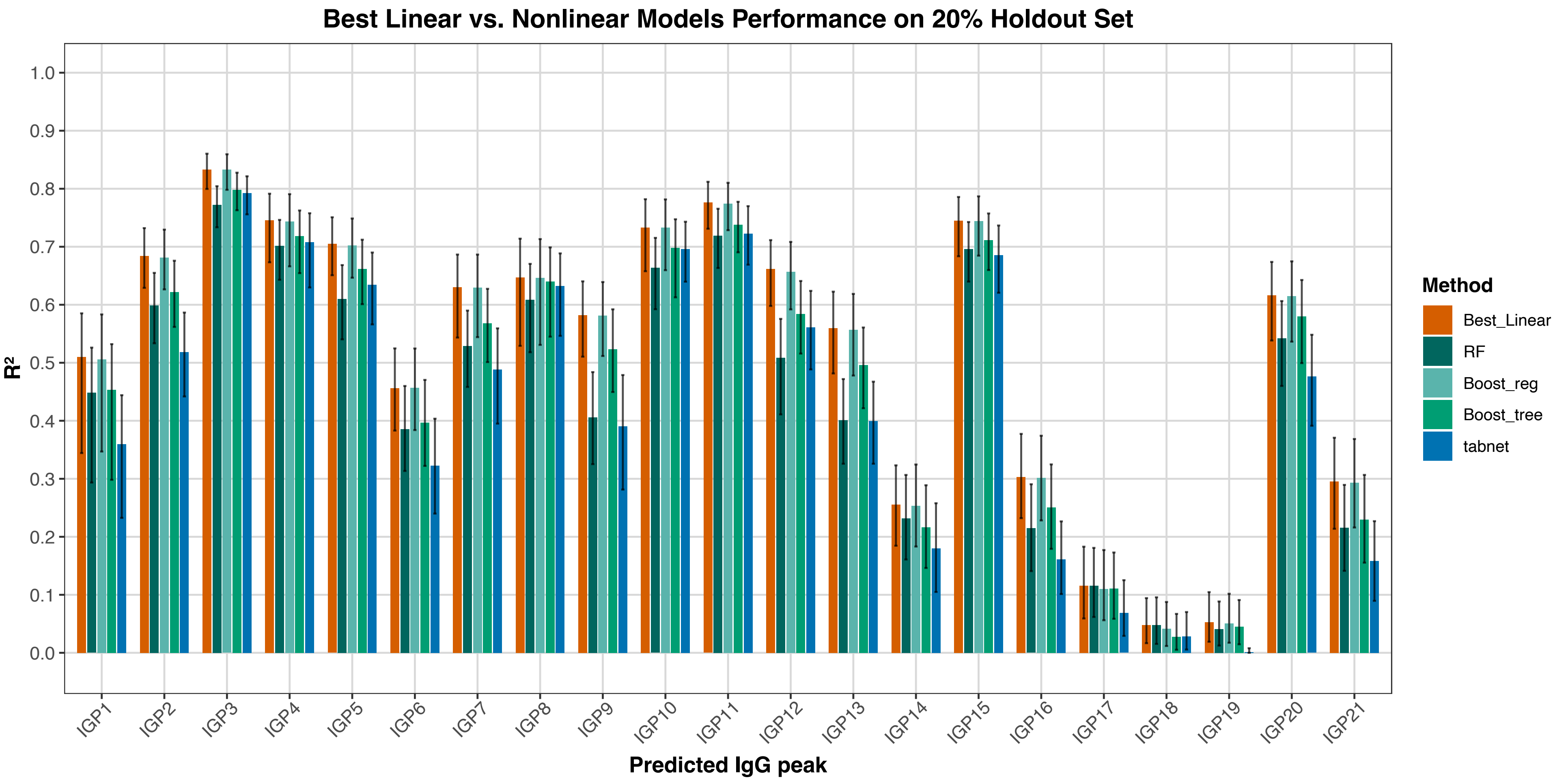


Supplementary Figure 2. Comparative performance of the best linear model versus nonlinear models in predicting IgG glycan peaks. The plot shows $R^2$ values on a 20% hold-out set from the TwinsUK cohort. Models evaluated include: the best linear model, random forest (RF), Gradient Boosting (Linreg Boost and Tree Boost), and a deep learning model (TabNet).

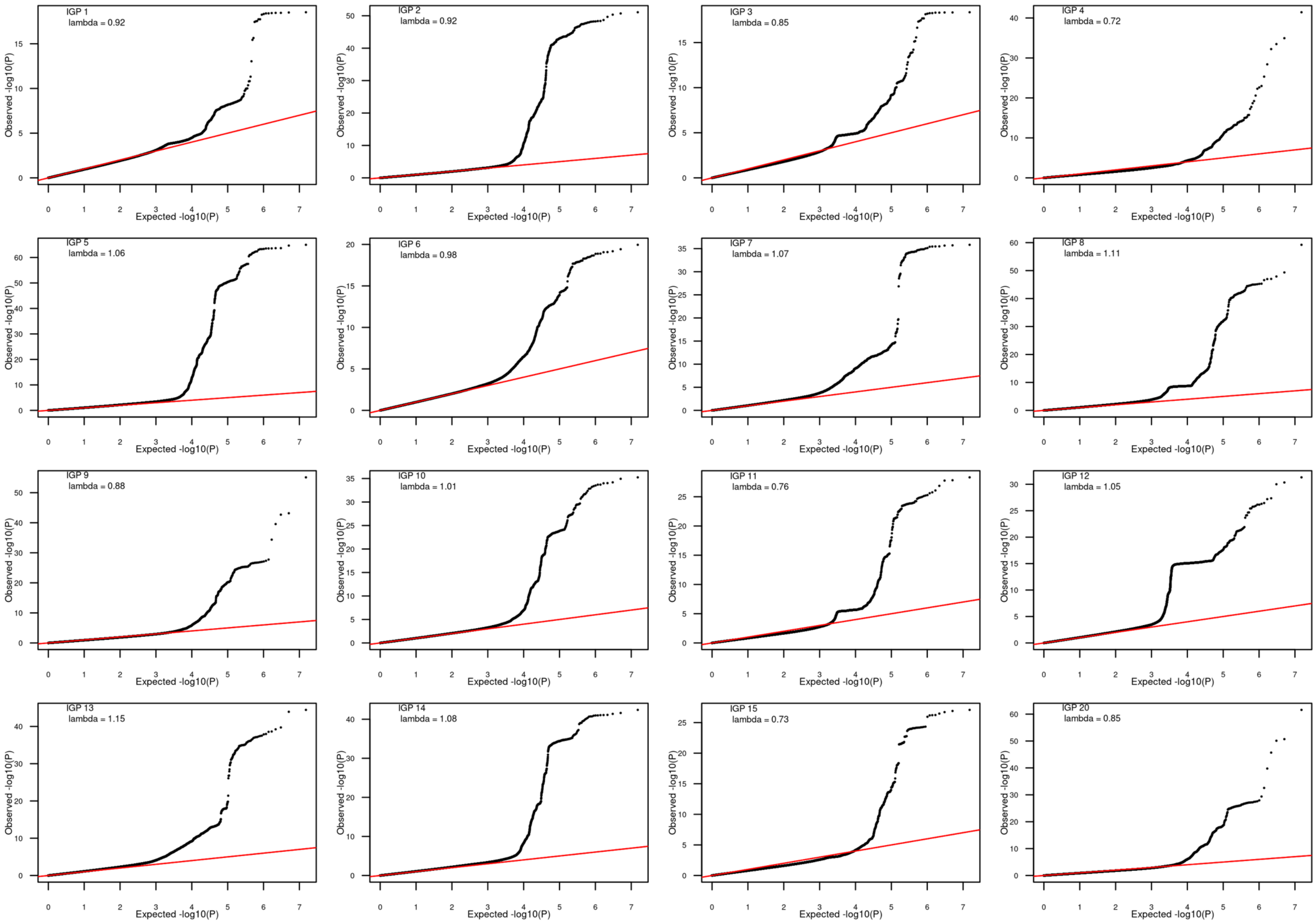
IGP 1
lambda = 0.92
IGP 2
lambda = 0.92
IGP 3
lambda = 0.85
IGP 4
lambda = 0.72
IGP 5
lambda = 1.06
IGP 6
lambda = 0.98
IGP 7
lambda = 1.07
IGP 8
lambda = 1.11
IGP 9
lambda = 0.88
IGP 10
lambda = 1.01
IGP 11
lambda = 0.76
IGP 12
lambda = 1.05
IGP 13
lambda = 1.15
IGP 14
lambda = 1.08
IGP 15
lambda = 0.73
IGP 20
lambda = 0.85
Observed -log10(P)
Expected -log10(P)

Supplementary Figure 3. QQ plots of GWAS association p-values for each IgG N-glycan peak. Sixteen panels show quantile-quantile plots comparing observed vs. expected $-\log_{10}$(p-values) for predicted IgG N-glycosylation GWAS. Each panel corresponds to one glycan peak. The genomic control inflation factor (lambda) is indicated within each plot.

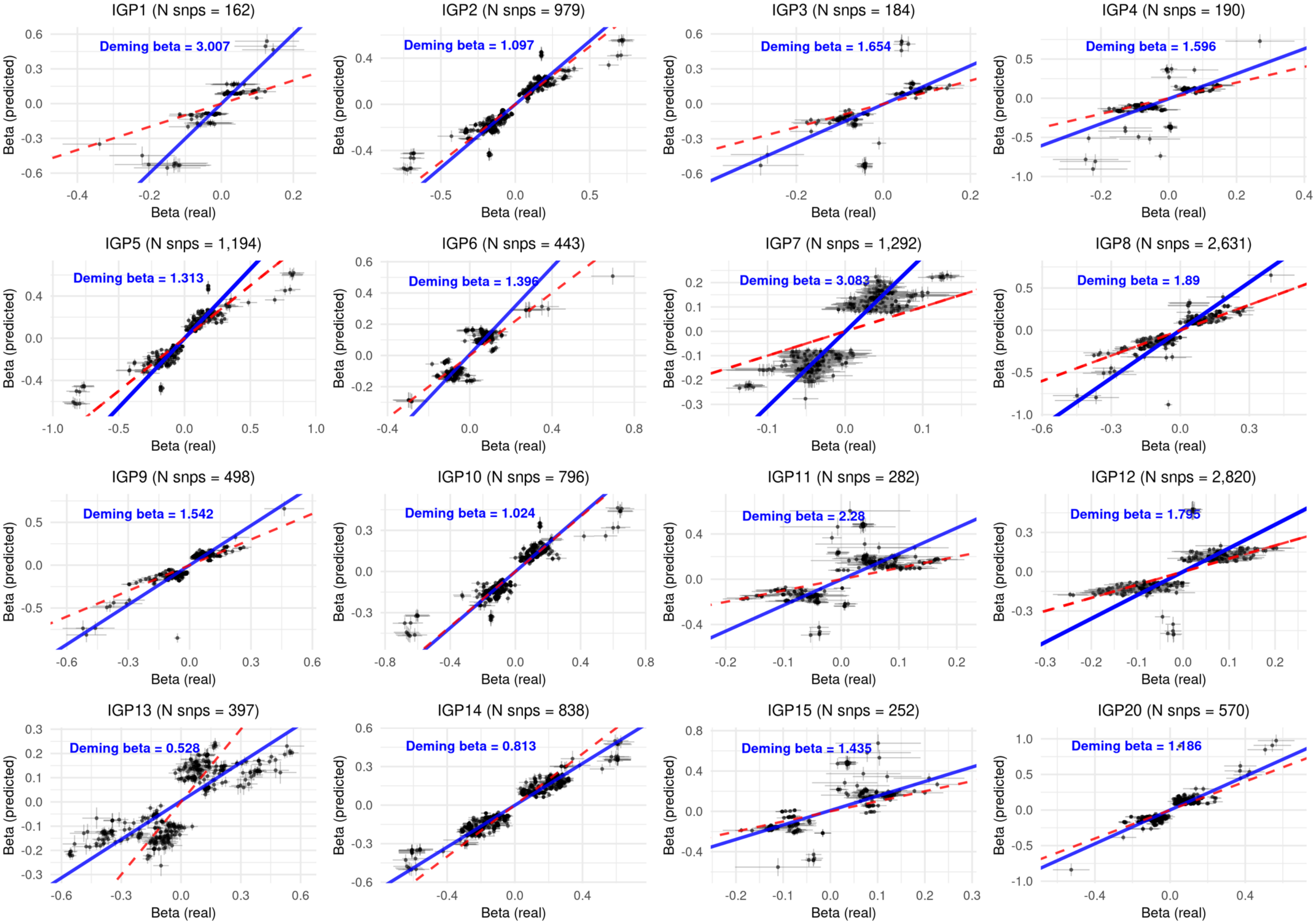

IGP1 (N snps = 162)
Deming beta = 3.007
IGP2 (N snps = 979)
Deming beta = 1.097
IGP3 (N snps = 184)
Deming beta = 1.654
IGP4 (N snps = 190)
Deming beta = 1.596
IGP5 (N snps = 1,194)
Deming beta = 1.313
IGP6 (N snps = 443)
Deming beta = 1.396
IGP7 (N snps = 1,292)
Deming beta = 3.083
IGP8 (N snps = 2,631)
Deming beta = 1.89
IGP9 (N snps = 498)
Deming beta = 1.542
IGP10 (N snps = 796)
Deming beta = 1.024
IGP11 (N snps = 282)
Deming beta = 2.28
IGP12 (N snps = 2,820)
Deming beta = 1.795
IGP13 (N snps = 397)
Deming beta = 0.528
IGP14 (N snps = 838)
Deming beta = 0.813
IGP15 (N snps = 252)
Deming beta = 1.435
IGP20 (N snps = 570)
Deming beta = 1.186
Beta (predicted)
Beta (real)

Supplementary Figure 4. Effect size concordance for genome-wide significant SNPs across IgG N-glycan peaks. Sixteen panels compare effect size estimates (beta ± standard error) from the observed IgG glycan GWAS (x-axis) versus the plasma-predicted IgG glycan GWAS (y-axis) for SNPs with genome-wide significance ($p < 5\times10^{-8}$) in at least one study. Blue lines show the Deming regression fit (slope Deming beta), which accounts for standard error in both variables; red dashed lines indicate y = x (perfect concordance).

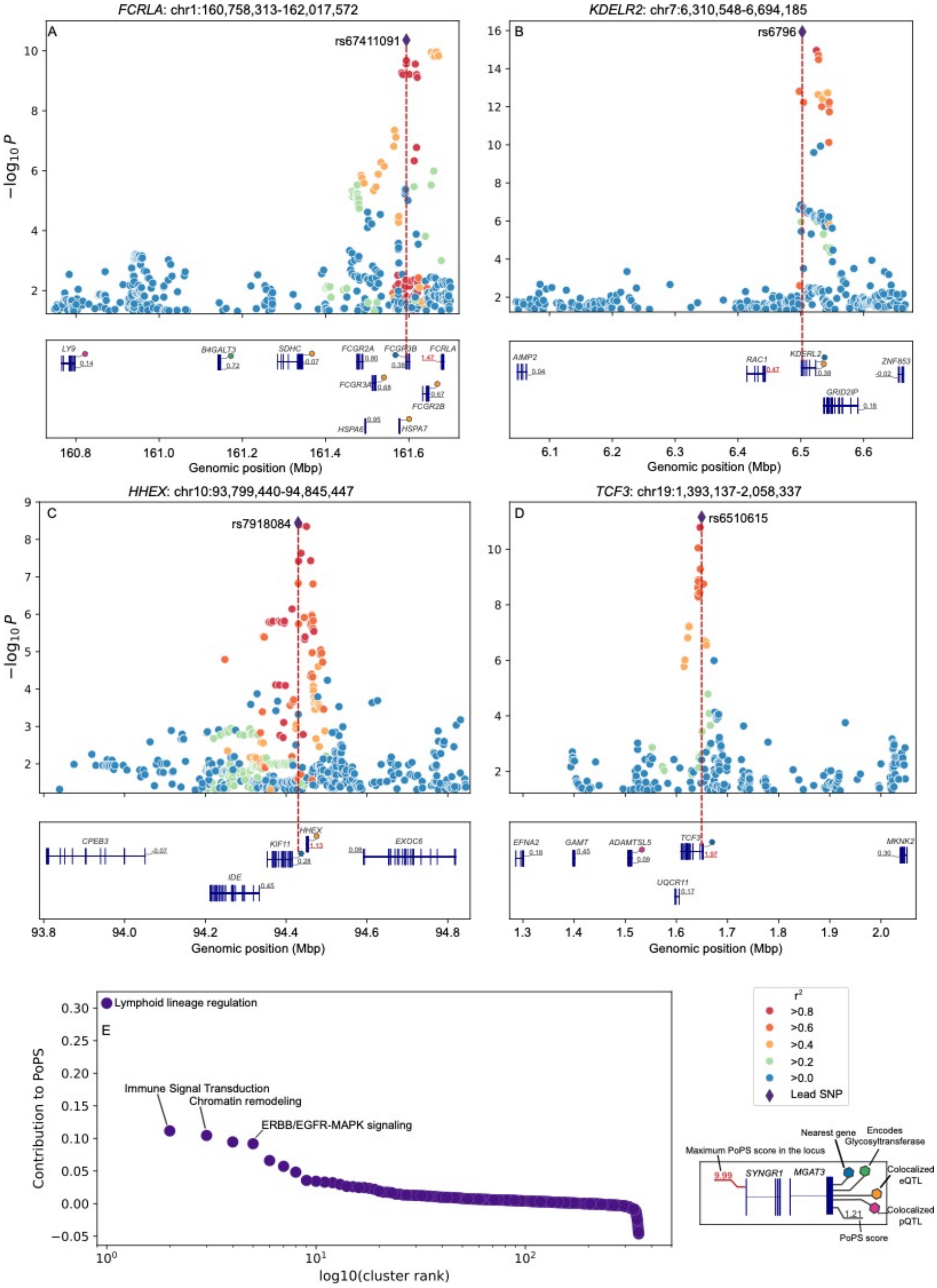

FCRLA: chr1:160,758,313-162,017,572
A
rs67411091
−log10 P
LY9
B4GALT3
SDHC
FCGR2A
FCGR3B
FCRLA
FCGR3A
FCGR2B
HSPA6
HSPA7
Genomic position (Mbp)
KDELR2: chr7:6,310,548-6,694,185
B
rs6796
AIMP2
RAC1
KDELR2
ZNF853
GRID2IP
Genomic position (Mbp)
HHEX: chr10:93,799,440-94,845,447
C
rs7918084
−log10 P
CPEB3
HHEX
KIF11
EXOC6
IDE
Genomic position (Mbp)
TCF3: chr19:1,393,137-2,058,337
D
rs6510615
EFNA2
GAMT
ADAMTSL5
TCF3
MKNK2
UQCR11
Genomic position (Mbp)
Lymphoid lineage regulation
E
Immune Signal Transduction
Chromatin remodeling
ERBB/EGFR-MAPK signaling
Contribution to PoPS
log10(cluster rank)
r²
>0.8
>0.6
>0.4
>0.2
>0.0
Lead SNP
Nearest gene
Encodes Glycosyltransferase
Maximum PoPS score in the locus
9.99
SYNGR1
MGAT3
Colocalized eQTL
Colocalized pQTL
1.21
PoPS score

Supplementary Figure 5. Candidate gene prioritization. A-E. Gene prioritization for loci tagged by rs67411091, rs6796, rs7918084, rs6510615. Regional association plots depict -$\log_{10}(P)$ for all markers in the vicinity of potential effector genes. Genes were considered potential effectors if prioritized by at least one of eight predictors: nearest gene to lead marker, direct experimental support linking the gene to N-glycosylation, known CDG gene, direct involvement in glycan biosynthesis, harboring damaging or coding variation, cis-eQTL or cis-pQTL colocalization with glyQTL in relevant tissues, or highest PoPS score in the locus. Genes ranking in the top five by PoPS are also shown. Colored hexagons denote prioritization criteria. Numbers indicate PoPS scores, with the locus maximum highlighted in red. E. Feature clusters contributing to causal gene prioritization. Rank-order plot of 348 feature clusters (arising from 5350 distinct features) contributing to the prioritization of likely causal genes for N-glycosylation by PoPS.
